# Effects of partner workload and increasing environmental temperature on nestling provisioning and body temperature in a declining aerial insectivore

**DOI:** 10.1101/2025.09.23.678107

**Authors:** Megan C. Heft, Bronwen Hennigar, Gary Burness

## Abstract

**Research Highlights:** In adult male tree swallows, we found that experimental manipulations of heat dissipation capacity had little effect on the provisioning rate, body temperature, nor growth of nestlings. That males did not respond to enhanced heat dissipation suggests that they may provision below the threshold of overheating. Rather, male provisioning was affected by the provisioning rate of the male’s partner and environmental temperature.

With climate change, birds will face increasing thermoregulatory demands, which may alter reproductive behaviors such as offspring provisioning. Experimental studies have shown that the provisioning capacity of female tree swallows (*Tachycineta bicolor*) is limited by their risk of overheating. Given that parental investment strategies may vary between sexes, the thermal environment may have a different impact on males. We experimentally trimmed ventral feathers from male tree swallows to create a “thermal window” through which they could dissipate heat. We remotely monitored provisioning rate and core body temperature of males and their female partners. At high temperatures, all males decreased their nestling provisioning rates irrespective of trimming treatment. In addition, trimmed males maintained core body temperatures similar to those of controls. This suggests that in contrast to previous work with females, males limit provisioning rates to levels below the threshold at which they would overheat. Regardless of male treatment, females adjusted their own activity to match that of their male partners; whether there are costs to females is unknown. Combined, these studies highlight that sex-specific differences in thermal physiology and behavior must be considered when predicting responses to climatic warming.

## Introduction

Climatic warming is predicted to result in an increase in extreme temperature and weather events (Sippel et al., 2020). In endotherms, prolonged exposure to high ambient temperatures has been associated with reduced reproductive success (e.g., Van de Ven et al., 2020; Pipoly et al., 2022, Levillan et al. 2025) and reports of mass mortality (Jones et al., 2018; Piatt et al., 2020; McKechnie et al., 2021). As thermal challenges are unlikely to subside given the current climatic estimates (IPCC, 2023), the extent to which individuals and populations will be able to effectively respond to thermal stressors is of growing concern (Altan et al., 2003; Heeter et al., 2023; Taff & Shipley, 2023).

Among endotherms, birds may be at particular risk of climatic warming given their relatively high body temperatures and elevated metabolic rates. When birds are faced with thermal stressors, they increase heat loss via evaporative or non-evaporative mechanisms (Tattersall et al., 2016) and/or allow body temperature (T_b_) to increase which may result in hyperthermia (e.g., Tapper et al., 2020b; Freeman et al., 2022). In species that exhibit extensive parental care, high environmental temperatures may exacerbate the effects of heat generated by parents during flight and in other activities associated with provisioning offspring (Tapper et al., 2020a). In fact, an individual’s rising body temperature and its inability to sufficiently dissipate metabolic heat generated during parental care may limit parental activity (heat-dissipation limit theory; Speakman & Król, 2010; Zagkle et al., 2022). The fitness benefits of increased provisioning rates must therefore be balanced against possible physiological damage induced by hyperthermia, including protein denaturation (Vazquez & Larson, 2013), depressed immune function (Alhenaky et al., 2017), and increased oxidative stress (Huang et al., 2015).

To date, a handful of avian field studies have explored experimentally how the risk of overheating impacts the capacity of individuals to maintain performance during a breeding event. For example, female tree swallows (*Tachycineta bicolor*) provided with an increased capacity to dissipate heat via experimental trimming of feathers, provisioned their nestlings at higher rates than their untrimmed counterparts, at least at highest temperatures (Tapper et al., 2020a). This suggested that at high environmental temperatures, an individual’s activity (and thereby capacity to care for young) was determined by a thermal constraint, as predicted by the heat-dissipation limit theory (Speakman & Król, 2010). In contrast, female blue tits (*Cyanistes caeruleus)* that were trimmed of feathers did not increase provisioning, but instead displayed enhanced immune function, suggesting a prioritization of self-maintenance (Nord & Nilsson, 2019; Andreasson et al., 2020).

An individual’s parental-investment strategy involves a trade-off between allocating energy and resources between current and future reproductive attempts (Clutton-Brock, 1991; Lemaître & Gaillard, 2017). In species that exhibit biparental care, thermal limits may play a role in pair coordination, with possible effects on investment strategies. Recently, it was shown that male tree swallows partnered with experimentally trimmed females increased their own provisioning efforts at high environmental temperatures, despite not being trimmed themselves (Tapper et al., 2020a). Male and female tree swallows are known the adjust their provisioning efforts based on partner quality (Lendvai et al., 2018). However, if males were able to increase their workload to better match that of their female partners, males were either working below their physiological limit, or perhaps willing to endure physiological damage (e.g., Dick & Guglielmo, 2019). In this sense, sex-specific investment strategies may emerge when males and females are faced with the removal of a possible physiological constraint (Nilsson & Nord, 2018).

Considering that reproductive strategies can vary between sexes (Kappeler et al., 2023), we investigated whether a male tree swallow’s capacity to dissipate heat would alter its reproductive investment, and we qualitatively compared our results with previous studies on females (Tapper et al 2020a, 2020b). We experimentally trimmed ventral feathers from male tree swallows to allow for enhanced heat loss and measured provisioning rates and body temperature in trimmed individuals and compared them with untrimmed controls. Further, we explored the response of their female partners (who were unmanipulated). We anticipated that the capacity to dissipate metabolic heat would limit activity in adult male tree swallows (i.e., the heat dissipation limit theory). We made the following predictions: (1) males with an augmented capacity to dissipate heat (trimmed males) would alter their reproductive investment, and increase their provisioning rate to nestlings across all environmental temperatures; (2) trimmed males would maintain a similar body temperature as untrimmed controls, by adjusting their activity to maintain T_b_ within a narrow range; (3) females paired with trimmed males would have higher nestling provisioning rates than females paired with non-trimmed males, because the provisioning rates of pairs are often coordinated (Lendvai et al., 2018); (4) nestlings from trimmed males would reach a heavier mass than nestlings from control males.

## Methods

### Study species and general field methods

All research was approved by the Trent University Animal Care Committee, in accordance with the Canadian Council on Animal Care (Animal Use Protocol # 26534). We conducted this study from May-July 2021, on two nest-box breeding populations of tree swallows located at the Trent University Nature Areas, Peterborough, Ontario, Canada (44°21′N, 78°17′W) and at the Lakefield Sewage Lagoon, Lakefield, Ontario (44°24’58.3“N 78°15’26.8”W). The Nature Areas site contains ∼40 boxes spread out in a grid across the site, while the Sewage Lagoon site consists of two large, central lagoons surrounded by a perimeter of ∼80 nest boxes. Further details on the sites are found in Fischer et al. (2020). At our study sites, tree swallows begin constructing their nests in early May, typically rearing 1 brood per season. Clutch size varies between 3-7 eggs, laid approximately one day apart. Incubation typically lasts 14 days with nestlings hatching relatively synchronously. After hatch, the nestling period spans approximately 15-22 days with adults providing food and maintenance care until the nestlings fledge (Winkler et al., 2020).

We monitored nests boxes daily from May through July to determine the lay and hatch dates for each clutch. As each egg was laid, we labelled it with a marker to establish order of laying and incubation dates. Incubation day 0 was the date that the last egg was laid, and hatch day 0 was the date the majority of nestlings in a brood hatched.

### Experimental manipulation of heat dissipation

When nestlings were between days 3-4 post-hatch, we captured adult male tree swallows and performed a feather-trimming manipulation. We assigned males randomly to either a Trimmed or Control treatment by flipping a coin (Trimmed: n=15, Control: n=20). Males in the Trimmed group had an approximately 3 x 2 cm area of ventral feathers trimmed with scissors, such that approximately 7% of body surface area was exposed, following Tapper et al. (2020a). To ensure the degree of trimming was consistent among individual males, we used a cardboard “stencil,” placed over each individual’s ventral surface. Males assigned to Control groups were handled for a similar length of time as the Trimmed individuals, but no feathers were removed.

Following trimming (or handling, in case of Controls), we implanted each male with either a temperature-sensitive passive integrated transponder (PIT) tag (accuracy ±0.5°C; Biotherm13; Biomark, Boise, Idaho, USA,) or attached a non-temperature-sensitive leg-mounted RFID tag (EM4100; #11001, GAO RFID, Ontario, 166 Canada). Tag type was randomized by coin flip. To implant a PIT tag, we first sterilized the incision site with ethanol, implanted the tag subcutaneously into the nape of the neck, and then sealed the incision site with tissue adhesive bond (Vetbond Tissue Adhesive, Fisher Scientific, Waltham, MA, USA). Prior to release we collected a 50-75 μL blood sample (as part of another study) and recorded body morphometrics (body mass, wing chord (flattened), and head-bill length). Handling time was < 10 minutes for each individual. Given limitations due to synchrony at hatch, loggers were rotated among nests and fewer birds were recorded than were initially trimmed and tagged. Sample size for temperature sensitive tagged males: Trimmed males: n= 4, Control males: n= 9. Sample size for RFID tagged males: Trimmed males: n= 9, Control males: n= 9.

To test female responsiveness to male trimming, we captured females at the same time as males and equipped each female with the same tag type as their male mates received (i.e, females paired with males that were equipped with temperature-sensitive tags were also assigned temperature-sensitive tags). We collected a blood sample and recorded the same morphometrics as for males; no females were trimmed (females paired with Trimmed males: n= 13; females paired with Control males: n= 17). Individuals were then released. One female paired with a Control male was tagged, but no nestbox visits were recorded while loggers were deployed. As a result, the sample size for females is slightly lower than that of males.

### Remote Monitoring

For all tagged adults we recorded nest-box visitation rates, while for individuals implanted with temperature-sensitive tags we also recorded subcutaneous body temperature (T_b_). We considered T_b_ to be a proxy for core body temperature (McCafferty et al., 2015; Andreasson et al., 2023). For individuals with temperature-sensitive tags, we recorded visitation rates and T_b_ using either a Biomark HPR Plus or a Biomark Small Scale Monitoring System (AR650) reader. Because of a limited number of Biomark readers, we cycled them among nests to capture adult visitation rates during 2-3 day windows throughout nestling development (early, middle, and late from 2-5 days, 6-9 days, and 10-14 days post-hatch, respectively). Readers (connected to 12v batteries) were placed at the base of each nest box and attached to a loop antenna mounted to the nest entrance. Visits were recorded anytime an individual entered the nest or perched at the nest entrance. Both types of Biomark readers were set to a 10 second ‘delay’ to limit tag detections when a bird spent a prolonged period in the nest box (Tapper at al., 2020). Individual reads included the date, time, and the bird identification code for a given tag (recorded when each tag was initially deployed).

To record visitation rates of individuals with leg mounted tags (rather than implanted temperature-sensitive tags), we used Generation 2 RFID readers (Cellular Tracking Technology, Rio Grande, NJ, USA) (Bonter and Bridge, 2011). Because we had a large number of Generation 2 RFID readers, we did not need to cycle them among nests. Nonetheless, to allow for comparison with data from temperature sensitive tags, we blocked our non-temperature sensitive detection data into 2-3 day windows to match the cycling of the Biomark readers.

### Environmental Data

We collected hourly readings for environmental temperature (°C), relative humidity (%), and windspeed (km ·hr-1), and total daily precipitation (mm) from the Trent University weather station (1.5 km from the Trent University Nature Areas and 9.5 km from Lakefield Sewage Lagoons). All environmental data were downloaded from Environment Canada (available at https://climate.weather.gc.ca/index_e.html).

### Nestling Morphometrics

Upon hatching of nestlings, we marked each nestling with a colored marker, allowing us to track the growth rate of individuals (until they were large enough to band). We weighed nestlings between the hours of ∼1200 – 1800 on days 0 (hatch), 3, 6, 9 and 12 post-hatch on a digital portable scale (Smart Weigh Digital Pro Pocket, ± 0.01 g). No measurements were recorded after day 12 to prevent premature fledge. In our analysis, nestlings were assigned to either a Control (n = 20) or Trimmed (n = 15) group based on the manipulation of the father, though no nestlings were manipulated. Note that the sample sizes for nests of nestlings were greater than those for adults used in provisioning rate analyses (above); due to logger limitations (cycling loggers between boxes) four nests could not be included. However, males were still captured and manipulated at these nests.

### Data compilation

#### Nest box visitation rate

All data organization and analysis was performed in R (R version 4.2.2, R Core Team). To quantify RFID reads (from both thermal and nonthermal tag types), we transformed raw RFID reads into nest box visits using the ‘visits’ function from feedr (LaZerte et al., 2017; Tapper et al., 2020a). We considered a nest box visit to be an index of parental provisioning. Subsequent statistical analyses for parental provisioning visits were then completed using these transformed data. Observations for adult provisioning visits were balanced within pairs (n=133 male observations, n=144 female observations).

Because males were captured between days 3-4 of nestling development, we restricted our analyses of provisioning visits to days 3-14 of nestling development. In addition, adults do not typically feed nestlings between the hours of ∼21:00 and 05:00 (Tapper et al., 2020a), so we excluded this window from our provisioning rate data. In the case where successive RFID reads occurred within 60 seconds (i.e., a bird was present in the box for multiple reads within the span of 60 seconds), we defined this as a singular visit. In our analysis, provisioning rate was defined as the average number of provisioning visits per hour of daytime activity (i.e., between 05:00 and 21:00 each day), or per hour that a logger was recording at a given nest box (if the logger was removed partway through the day).

### Statistical analysis

#### Provisioning rate

For our analyses of male and female provisioning rates, and male body temperature we evaluated competing models with an Akaike Information Criterion corrected for small sample sizes (AICc). In our analysis, we considered models with a ΔAICc of < 2 to be strongly supported (Burnham & Anderson, 2004). For our provisioning rate and body temperature models, we evaluated models based on model weights and likelihoods (Hurvich & Tsai, 1989).

For each of our feeding rate and body temperature analyses, we fit a series of candidate models and report the number evaluated: male provisioning (n = 16), female provisioning (n = 16), and male body temperature (n = 10). We also report a null model (intercept only) for reference. For models with strong support, we report the 95% confidence intervals (CI) and associated effect size for individual terms.

#### Statement on AI

Artificial Intelligence (AI) was used during data analysis to refine visualizations and to suggest corrections to code. Specifically, ChatGPT (OpenAI, 2025) was used to improve layouts, color scheme, and labeling of figures, for clearer visualizations. All code was reviewed and checked for accuracy.

#### Male provisioning rate

In our analysis of male provisioning rate, we were primarily interested in the response of males to our experimental manipulation (trimming). To explore potential factors contributing to variation in average number of visits per hour, we evaluated combinations of predictors including manipulation (trimmed or control, i.e., untrimmed), tag type (thermal or nonthermal), brood size (at hatch), clutch initiation date, average daily environmental temperature, relative humidity and wind speed, and total daily precipitation. We evaluated environmental predictors for collinearity by running a variance inflation factor test (all VIF <3; ‘vif’ function, Car package, version 3.1-2; Fox and Weisberg, 2019). We also included the average number of visits per hour for females (filtered from 05:00 to 21:00). Based on the nonlinear relation between male provisioning rate and environmental temperature, we included a quadratic term for average daily environmental temperature (T_a_^2^). Since previous work has shown that the effects of treatment may vary based on environmental temperature (Tapper et al., 2020a), we also tested for an interaction between average daily environmental temperature and manipulation (along with an interaction between manipulation and the quadratic; T_a_^2^). To account for repeated measures from the same individual over the brood rearing period, we included a random intercept for individual male’s identity.

Since thermal loggers were cycled among nests (sometimes being moved partway through the morning) we favored an averaging approach for provisioning rates, where feeding rate is defined as the total number of visits per day divided by the total number of hours logged per day. As in previous work (Tapper et al., 2020a), the cycling of thermal loggers constained our analysis of adult provisioning as rotations led to unequal observation effort at nest boxes of birds equipped with thermal tags. While an hourly analysis was considered, we preferred the average hourly provisioning rate, to account for unequal coverage and allow for comparisons between tag types. However, we also conducted additional analysis of raw hourly provisioning with focal hours captured using hourly provisioning rate and hourly environmental variables (with the same models sets as described above for average hourly, to consider finer scale hourly variation throughout the day). We provide the output of these analyses in supplementary table S1 and S2.

#### Male body temperature, T_b_

We tested the impact of experimental trimming on male average hourly T_b_ (°C), with nestling age, average number of visits per hour, manipulation (trimmed or untrimmed), and hour of the day (filtered for diurnal readings; between 05:00 and 21:00), average daily environmental temperature, relative humidity, and wind speed as main effects. As the relation between environmental temperature (T_a_) and body temperature (T_b_) was not linear, we also tested temperature as a linear (T_a_), quadratic (T_a_^2^) and cubic (T_a_^3^) term. We detected five T_b_ observations that had standardized residuals > 3.0, but since the values were within the range reportedly previously (Tapper et al 2020b), we retained them in our analysis. To account for repeated measures of T_b,_ we included a random intercept for individual male’s identity.

#### Female provisioning rate

In addition to understanding factors that contribute to variation in male provisioning rate, we were also interested in how a male’s behavior, and particularly the experimental trimming treatment, may impact that of the female partner. Due to heteroscedasticity and non-normality of residuals in the female provisioning rate model, we applied a natural logarithmic transformation (ln) to the response term (average number of visits per hour). Where relevant, we back-transformed values and confidence intervals to the original scale. In our analyses of female provisioning rate, we included the partner’s manipulation (Trimmed or Control (ie., untrimmed)), brood size at hatch, clutch initiation date, and average daily environmental temperature, relative humidity and wind speed, and total daily precipitation as main effects. We also tested an interaction term of average daily environmental temperature and male manipulation, since previous work suggested that untrimmed partners may respond to a partner’s trimming manipulation (Tapper et al., 2020a). To account for female identity, we included a random intercept for individual bird identity. As with males, we report the average provisioning rate for females (total visits per day divided by total hour recorded per day), but female provisioning rate was also analyzed with hourly provisioning rate and environmental variables (see supplementary materials). Note: We did not report T_b_ of adult females as female T_b_ has previously been reported on in (Tapper et al., 2020b).

#### Nestling growth rate

We calculated nestling growth rate using a three-parameter logistic regression model, following Tapper et al., (2020a). The logistic growth function models the sigmoidal increase in nestling mass with age. For the nestling growth rate model, we calculated the asymptote (*A*; the upper limit of the growth curve), the inflection point (*I*; the point at which the growth rate is greatest), and growth rate constant (*K;* the steepness of the growth rate curve). For starting estimates in our growth curves, we used the “SSlogis” function (Stats). We include the five measurements per nestling (days 0, 3, 6, 9, and 12 post-hatch), which encompasses all major phases of the growth curve (early, middle, and late; McCarty, 2001) and provides sufficient information for estimates.

Growth curves were fitted in a single hierarchical model for all nestlings, rather than fitting individual treatment groups or individuals usings a nonlinear mixed-effect structure. We account for the lack of independence between nestlings from the same brood with box identity as a random effect. Similarly, to account for repeated measures from the same nestling, we also included nestling identity as a random effect. Since individual variation can differ at different stages of growth, we compared three growth models, each with different random effects structures based on the three growth parameters (*A, I, K*). Each model included nestling and box identity as random effects, but in each model we allowed for variance in one growth parameter while holding the other two constant. All models included the same fixed effects and were assessed for different random effect structures using maximum likelihood, to determine which random effect structure best explained variance in the different growth curves. The model with the highest log-likelihood was considered best supported. We included brood size at hatch as a covariate to account for possible variation between large and small broods.

## Results

### Male provisioning rate was best predicted by environmental temperature and the provisioning rate of female partners

Variation in average hourly provisioning rate by males was best predicted by environmental temperature (T_a_), its quadratic term (T_a_^2^), and average hourly female provisioning rate, each of which appeared in our only model with strong support (ΔAIC_C_ < 2.0; Table 1). Our top model was >10^5^ times more likely to be the top model than the null (Acc W_i_ = 1.0; ER = 1.0, Table 1), we therefore report parameter estimates and effects from this model alone. As environmental temperature (T_a_) increased up to approximately 20°C, males increased their provisioning rate (T_a_; = 3.678); although provisioning rates decreased at highest environmental temperatures (T_a_^2^; = −0.099; Fig 1; Table 2). Within a breeding pair, male provisioning rate increased with increasing female provisioning rate ( = 0.482; Table 2). Contrary to our prediction – that trimmed males would increase provisioning rates across all environmental temperatures, the trimming manipulation was not a strong predictor of male provisioning rate. That is, while manipulation appeared in our top model (ΔAIC_C_ < 2.0; Table 1), the confidence intervals crossed zero (Table 2). On average, male provisioning rate (visits·hr^-1^) did not differ between treatments (LSM ± SE: trimmed birds: 10.6 ± 0.50; control birds: 10.9 ± 0.40). Likewise, while we tested whether the effects of male trimming were dependent on environmental temperature - as previous work suggested that trimmed birds may be able to work harder at high temperatures (Tapper et al., 2020a) - neither the interaction term of the first nor second order polynomial (manipulation × T_a_; manipulation × T_a_^2^) appeared in any of our top models. While hourly data were also analyzed, the general patterns of male provisioning were consistent between both temporal scales, with female provisioning rate appearing as a strong predictor at both the average hourly (Table 1) and hourly scales (Table S1).

**Figure 1.**
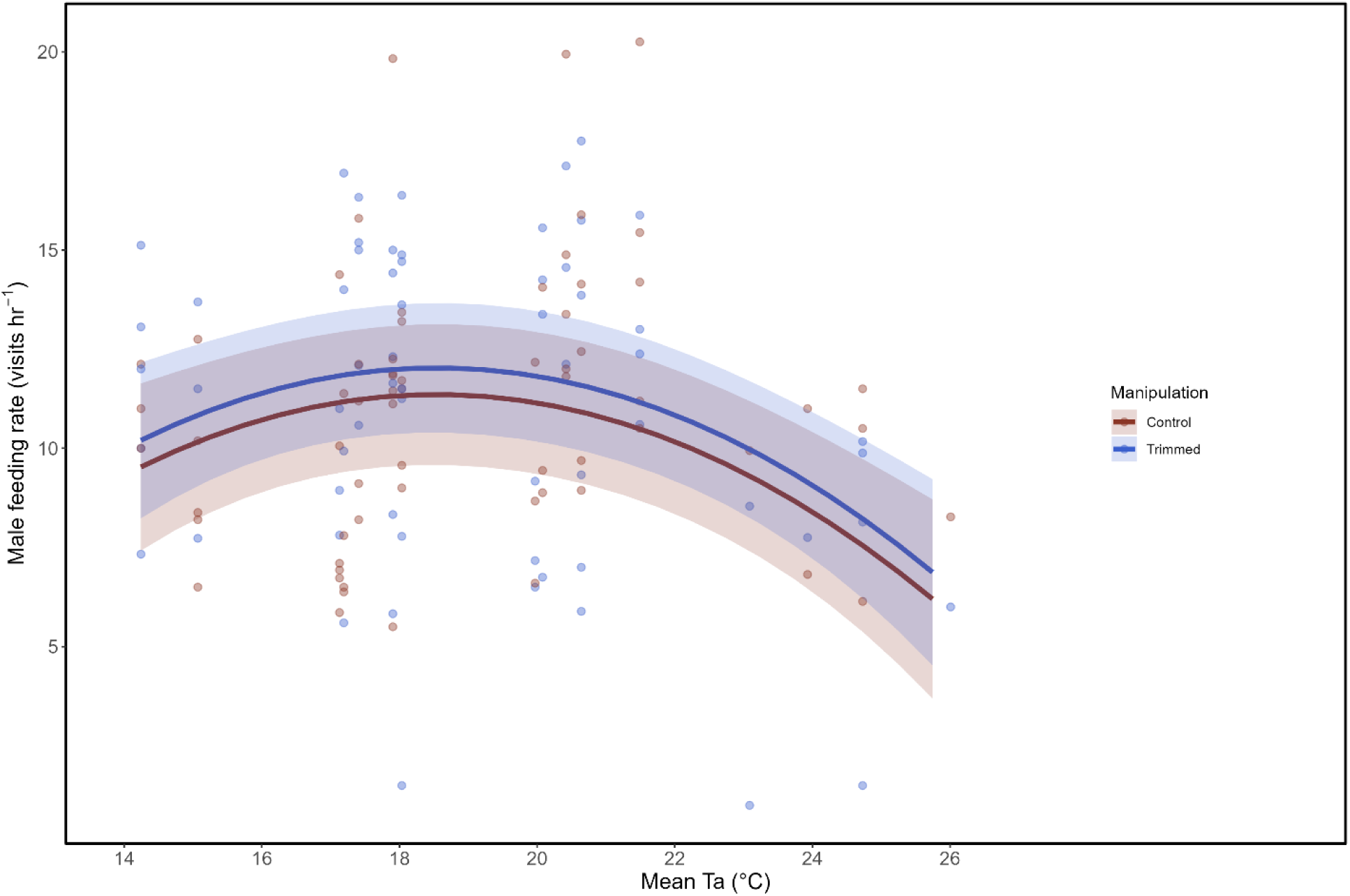
Male average hourly provisioning rate decreased at high temperature (T_a_), but did not differ between trimmed (blue) and control (red) males. Figure is drawn from the best-supported model, with all other fixed effect values held at their means. Bands and trend lines indicate the predicted means and 95% CI. Individual data points represent average provisioning rate for each individual bird for a given temperature (Trimmed: n=13, Control: n=18).

**Table 1.** Model selection results for factors that explain variation in average hourly provisioning rate by adult male tree swallows. Of the 16 models run, there was only a single model with strong support (<2 ΔAIC_C_). Models were run with a random effect for individual bird identity (Trimmed: n=13, Control: n=18).

| Models | K | AIC<sub>C</sub> | $\Delta AIC_C$ | W<sub>i</sub> | Acc W<sub>i</sub> | R<sup>2</sup><sub>m</sub> | ER |
| --- | --- | --- | --- | --- | --- | --- | --- |
| T<sub>a</sub> + T<sub>a</sub><sup>2</sup> + Female Provisioning Rate + Manipulation | 6 | 629.81 | 0.00 | 1.0 | 1.0 | 0.735 | 1.00 |
| Null Model | 2 | 702.84 | 73.04 | 0.00 | 1.0 | 0.487 | $>10^5$ |

Note: K = number of parameters, AIC<sub>C</sub> = corrected Akaike Information Criterion, $\Delta AIC_C$ = difference of focal model AIC<sub>C</sub> compared to the lowest model AIC<sub>C</sub>; W<sub>i</sub> = Akaike weight; Acc. W<sub>i</sub> = Accumulated weight of models; R<sup>2</sup><sub>m</sub> = Marginal R<sup>2</sup>; ER = Evidence ratio; T<sub>a</sub> = Average daily temperature; T<sub>a</sub><sup>2</sup> = Quadratic term for average daily temperature; Female Provisioning Rate = Average hourly provisioning rate of females paired with males; Manipulation = Male manipulation (trimmed or untrimmed [Control]). The null model was included for reference but did not have strong support ( $\Delta AIC_C > 2$ ).

**Table 2.** Factors affecting the average hourly provisioning rate of adult male tree swallows. Estimates of fixed effects with 95% confidence intervals (CI) and *P*-values (bold signifies statistical significance) are shown. Models were run with a random effect for individual bird identity (Trimmed: n=13, Control: n=18).

| Predictor | Estimate | 95% CI | P-value |
| --- | --- | --- | --- |
| Intercept [Control] | -29.093 | -46.59, -11.60 | **0.001** |
| $T_a$ | 3.678 | 1.88, 5.47 | **<0.001** |
| $T_a^2$ | -0.099 | -0.15, -0.05 | **<0.001** |
| Female Provisioning Rate | 0.482 | 0.36, 0.61 | **<0.001** |
| Manipulation [Trimmed] | 0.673 | -1.84, 3.18 | 0.584 |

### Male body temperature did not differ with experimental trimming, but increased with environmental temperature and provisioning rate

We predicted that trimming manipulations would allow males to increase provisioning rates and maintain a similar T_b_ as controls, because of their increased surface area to dissipate heat. The top model explaining variation in male T_b_ was modeled with the linear, quadratic, and cubic term for environmental temperature, along with manipulation and male provisioning rate (ΔAIC_C_ < 2.0; Table 3). We found manipulation was not a strong predictor of body temperature (P = 0.23, Table 4; Fig 2). Rather, male T_b_ increased with temperature (*T*_a_) and its cubic term (*T*_a_^3^; both P <0.001; Table 4, Fig 2). Similarly, male T_b_ increased with male provisioning rate (P<0.001; Table 3). Male T_b_ was not significantly influenced by the quadratic term for temperature (*T*_a_^2^; P = 0.36; Table 4), despite being retained in the top model. Note: Because sample size was limited for trimmed males, these results must be interpreted with caution as the small sample size may limit power to detect treatment effects.

**Figure 2.**
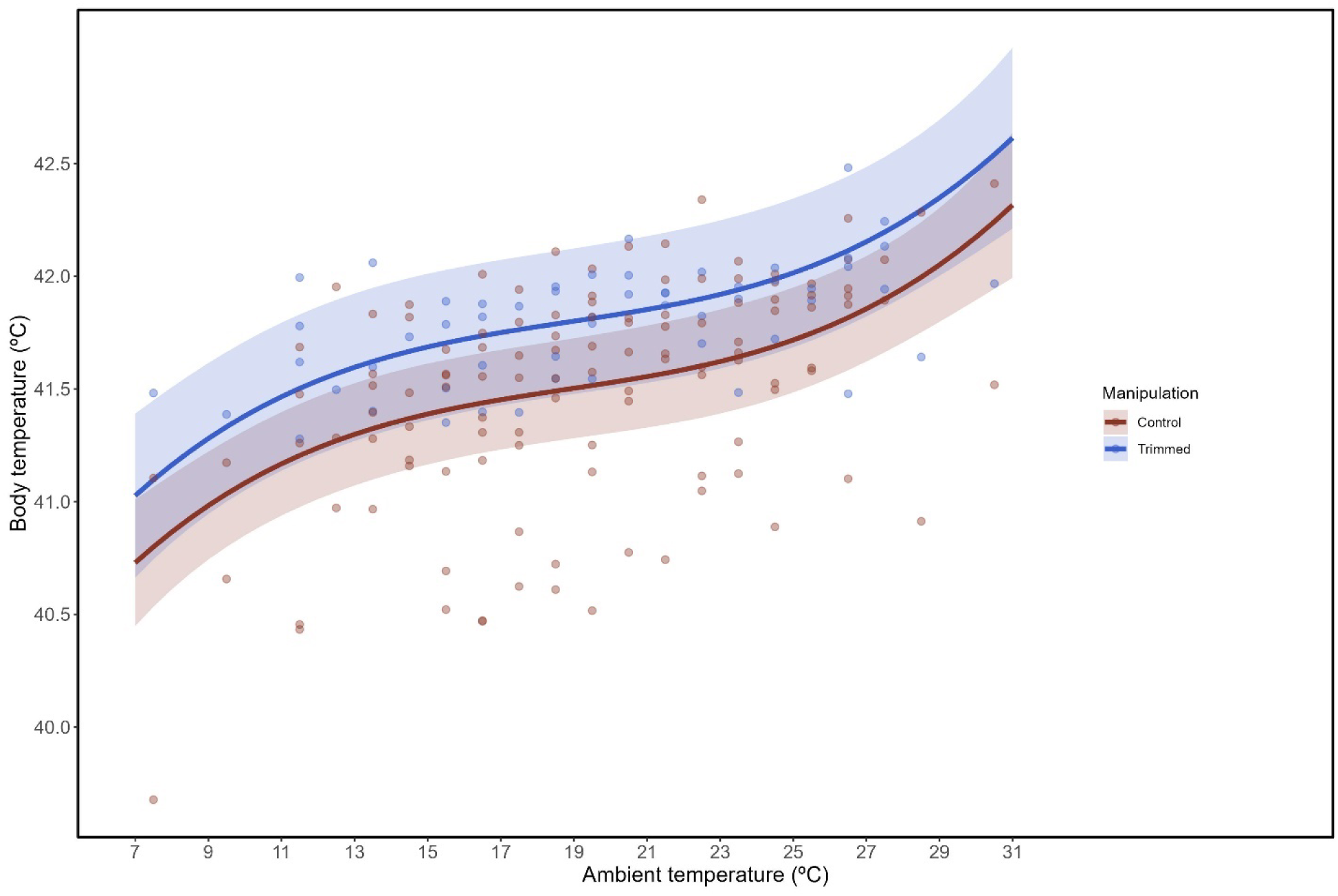
Male hourly body temperature increased with ambient temperature but did not differ between trimmed (blue) and control (red) males. Bands and trend lines indicate the predicted means and 95% CI. Individual data points represent average hourly T_b_ for each individual bird at a given temperature (Trimmed, n= 4; Control, n= 9). Note, this figure displays only the subset of males with temperature-sensitive PIT tags.

**Table 3.** Model selection results for factors that explain variation in the body temperature of adult male tree swallows. Of the 10 models run, there was only a single model with strong support (<2 ΔAIC_C_). Models were run with a random effect for individual bird identity (Trimmed, n= 4; Control, n= 9).

| Models | K | AICc | $\Delta AICc$ | Wi | Acc Wi | $R^2_m$ | ER |
| --- | --- | --- | --- | --- | --- | --- | --- |
| $T_a + T_a^2 + T_a^3 +$<br>Manipulation + Male<br>Provisioning Rate | 11 | -21.41 | **0.00** | 1.00 | 1.00 | 0.091 | 1.00 |
| Null Model | 4 | 95.18 | 116.59 | 0.00 | 1.00 | 0.00 | $>10^5$ |

Note: The null model was included for reference but did not have strong support ( $\Delta AIC_C > 2$ ).
$T_a^3$ = Cubic term for average daily temperature, all other abbreviations are as in Table 1

**Table 4.** Factors affecting the body temperature of adult male tree swallows. Estimates of fixed effects with 95% confidence intervals (CI) and *P*-values are shown. Bold indicates statistical significance. Models were run with a random effect for individual bird identity (Trimmed, n= 4; Control, n= 9). Approximately 12% of variance in body temperature was accounted for by individual identity.

| Predictor | Estimate | 95% CI | P-value |
| --- | --- | --- | --- |
| Intercept [Control] | 41.46 | 41.245, 41.674 | **<0.001** |
| $T_a$ | 3.55 | 2.885, 4.222 | **<0.001** |
| $T_a^2$ | 0.27 | -0.314, 0.863 | 0.363 |
| $T_a^3$ | 1.06 | 0.538, 1.598 | **<0.001** |
| Manipulation [Trimmed] | 0.24 | -0.170, 0.657 | 0.226 |
| Male Provisioning Rate | 0.01 | 0.004, 0.012 | **<0.001** |

### Female provisioning rate was best predicted by environmental temperature and the provisioning rate of male partners

The top model explaining variation in female provisioning rate included average daily environmental temperature (T_a_), male manipulation, and male provisioning rate (ΔAIC_C_ < 2.0; Table 5). There was no other competing model, and our top model was >10^5^ times more likely to be the top model than the null (Acc Wi = 1.0; ER = 1.0, Table 5), and we report parameter estimates and effects from this model alone. Females slightly increased their provisioning rate as environmental temperature increased ( = 0.02, Table 6; Fig 3). There was a small increase in female provisioning rate as male provisioning rate increased ( = 0.05, Table 6). However, while experimental manipulation of male partners appeared in our top model, the confidence intervals crossed zero, indicating that females paired with trimmed males did not adjust their provisioning rate based on male manipulation ( = −0.11, Table 6). Overall, female provisioning rate (visits·hr^-1^) did not differ with male manipulation (LSM ± SE: females partnered with trimmed males: 12.5 ± 0.57; females partnered with control males 13.6 ± 0.46). As with male provisioning rate, hourly data for females were also analyzed, with male provisioning rate and environmental temperature in top models for both temporal scales (Table 5; Table S2).

**Figure 3.**
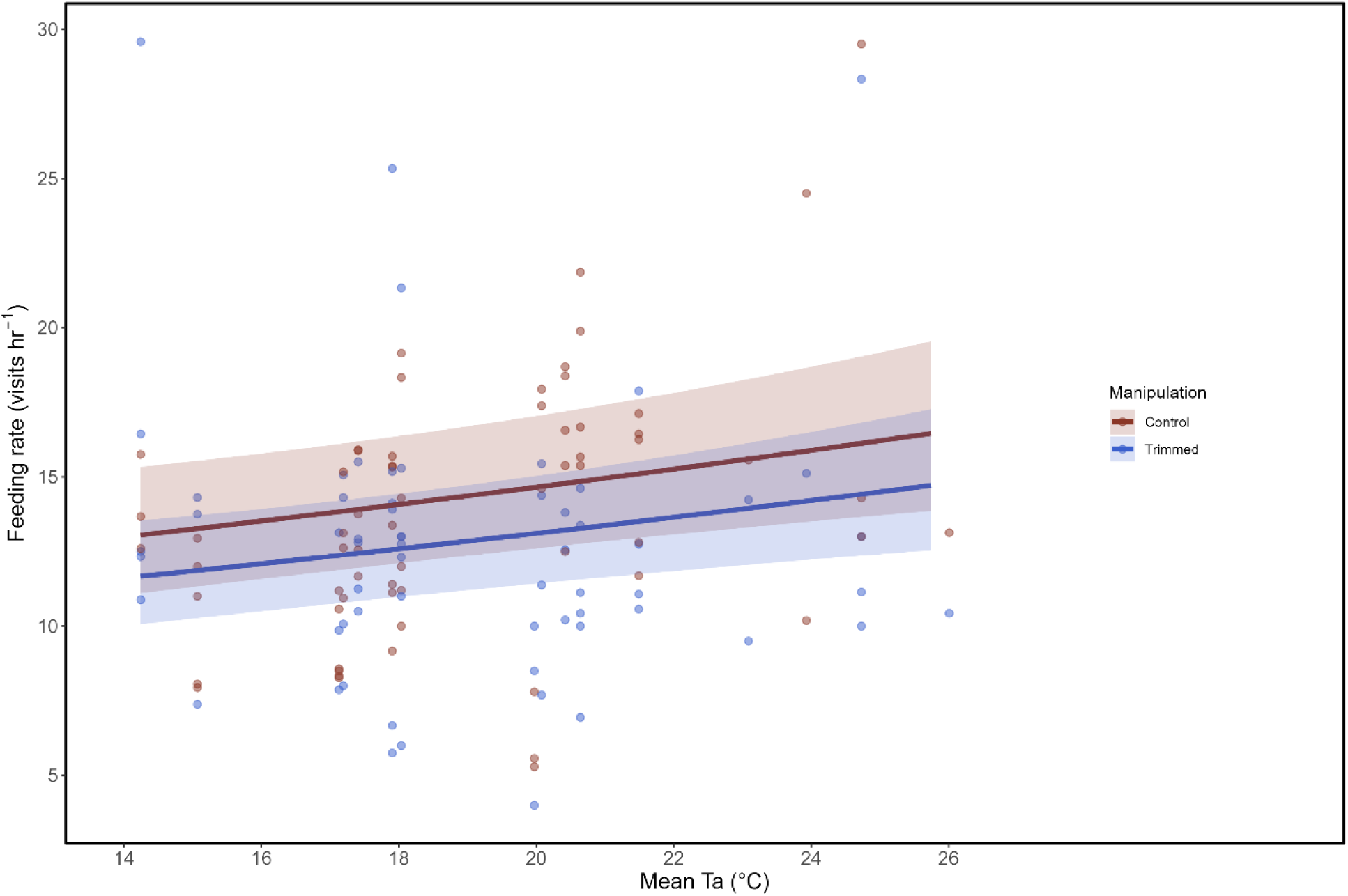
Females increased provisioning rates with increasing temperature (T_a_), but females partnered with trimmed males did not provision at a higher rate than those paired with control males. Sample sizes: females partnered with Trimmed males, n= 13; females partnered with Control males, n= 17.

**Table 5.** Model selection results for factors that explain variation in average hourly provisioning rate of adult female tree swallows paired with trimmed or untrimmed males. There was only a single model with strong support (<2 ΔAIC_C_). A total of 16 candidate models were run with a random effect for individual bird identity (females partnered with Trimmed males, n= 13; females partnered with Control males, n= 17).

| Models | K | AICc | $\Delta AICc$ | Wi | Acc Wi | $R^2_m$ | ER |
| --- | --- | --- | --- | --- | --- | --- | --- |
| $T_a$ + Manipulation + Male<br>Provisioning Rate | 5 | 11.07 | 0.00 | 1.00 | 1.00 | 0.765 | 1.00 |
| Null Model | 2 | 85.71 | 74.64 | 0.00 | 1.00 | 0.399 | $>10^5$ |

Note: The null model was included for reference but did not have strong support ( $\Delta AIC_C > 2$ ).
Abbreviations are as in Table 1

**Table 6.**
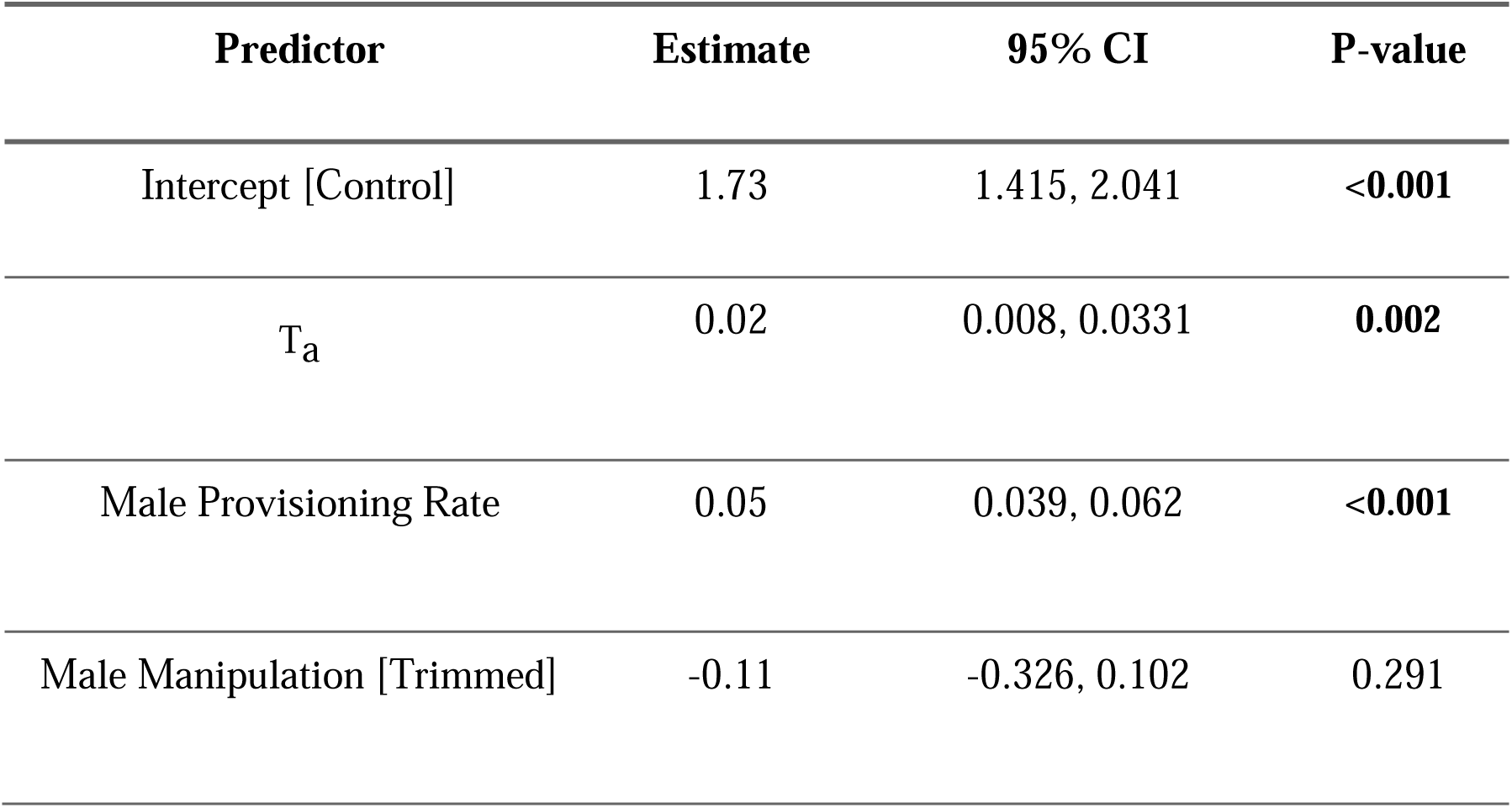
Factors affecting the average hourly provisioning rate of adult female tree swallows. Estimates of fixed effects with 95% confidence intervals (CI) and *P*-values are shown. Bold indicates statistical significance. Models were run with a random effect for individual bird identity (females partnered with Trimmed males, n= 13; females partnered with Control males, 858 n= 17).

| Predictor | Estimate | 95% CI | P-value |
| --- | --- | --- | --- |
| Intercept [Control] | 1.73 | 1.415, 2.041 | **<0.001** |
| T<sub>a</sub> | 0.02 | 0.008, 0.0331 | **0.002** |
| Male Provisioning Rate | 0.05 | 0.039, 0.062 | **<0.001** |
| Male Manipulation [Trimmed] | -0.11 | -0.326, 0.102 | 0.291 |

### Male trimming had a modest effect on nestling growth

Nestling growth rate was best modeled with a random effect structure for asymptotic mass (over models with a random intercept for the inflection point or growth constant; logLik = −2002.56). We found that male trimming (manipulation) did not affect the asymptote (*A*) of nestling mass (P = 0.471, Table 7). On average, nestlings from trimmed males weighed 22.3 ± 1.43g (± SE) at their asymptote (∼ day 12 post-hatch) while nestlings from control males were similar at 21.8 ± 1.25g (± SE; Fig. 4; Fig. 5). However, we found that nestlings from trimmed males developed slightly faster, reaching the inflection point of the growth curve (*I*) at 4.51 ± 0.25 days of age (± SE) compared to controls at 4.29 ± 0.23 days of age (± SE, P = 0.014, Table 7). We also found that nestlings from trimmed males had a slightly steeper curve for growth rate (i.e., growth rate constant) than controls (*K;* P = 0.005, Table 7), growing faster after reaching the inflection point of growth (nestlings from trimmed males: 2.10 ± 0.21; control nestlings: 1.88 ± 0.19; ± SE).

**Figure 4.**
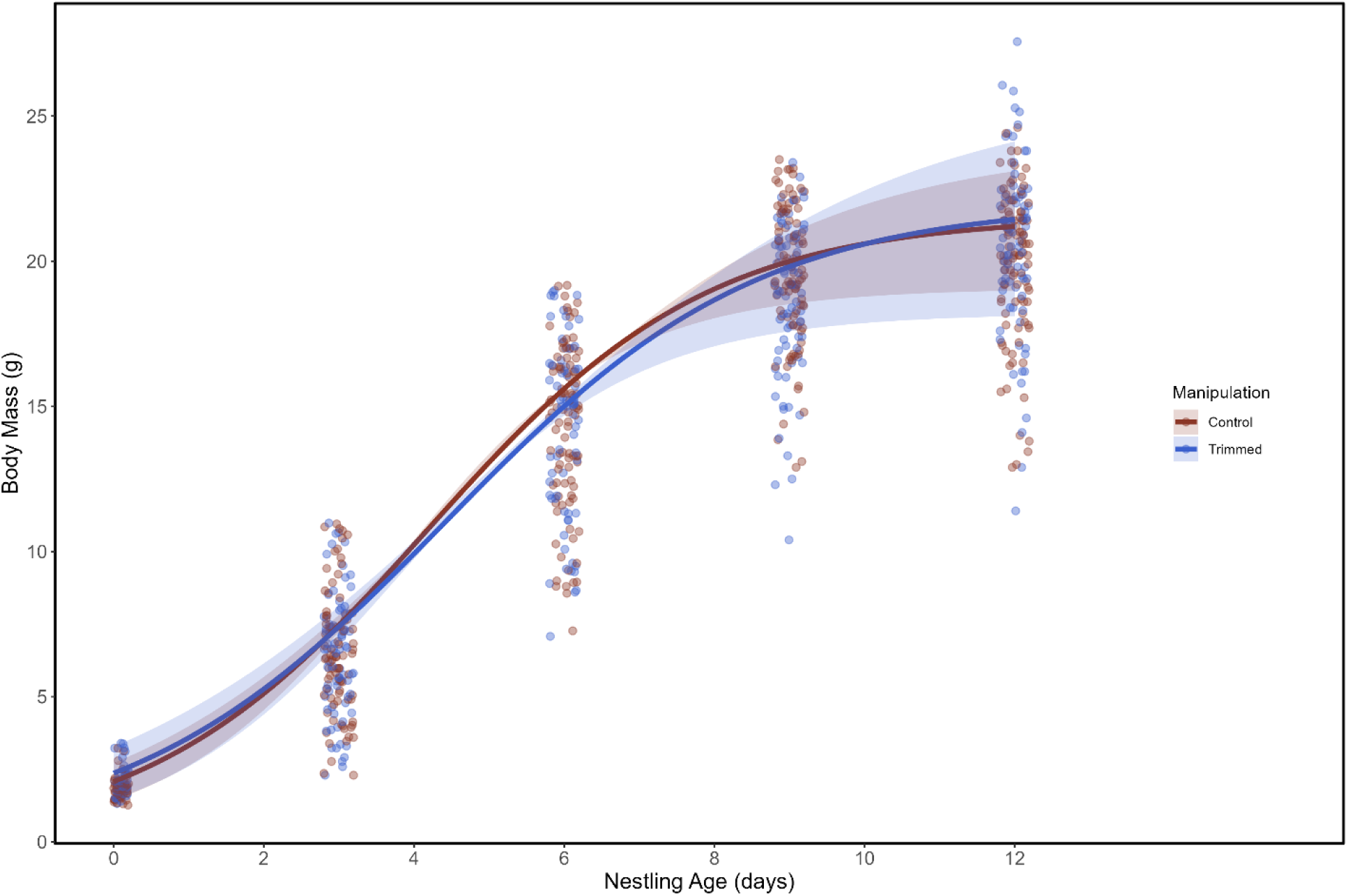
Estimated growth trajectories for nestlings from control and trimmed males. Nestlings from trimmed males reached a similar asymptotic mass as nestlings from control males. Growth estimates were based on a logistic growth curve using scale, midpoint, and asymptotes with random effects of nestling and box identity. Bands represent 95% CI. Each point represents the mass of a nestling on a given developmental day. Sample sizes: Control, n = 20 nests, n= 100 nestlings; trimmed, n = 15 nests, n= 86 nestlings).

**Figure 5.**
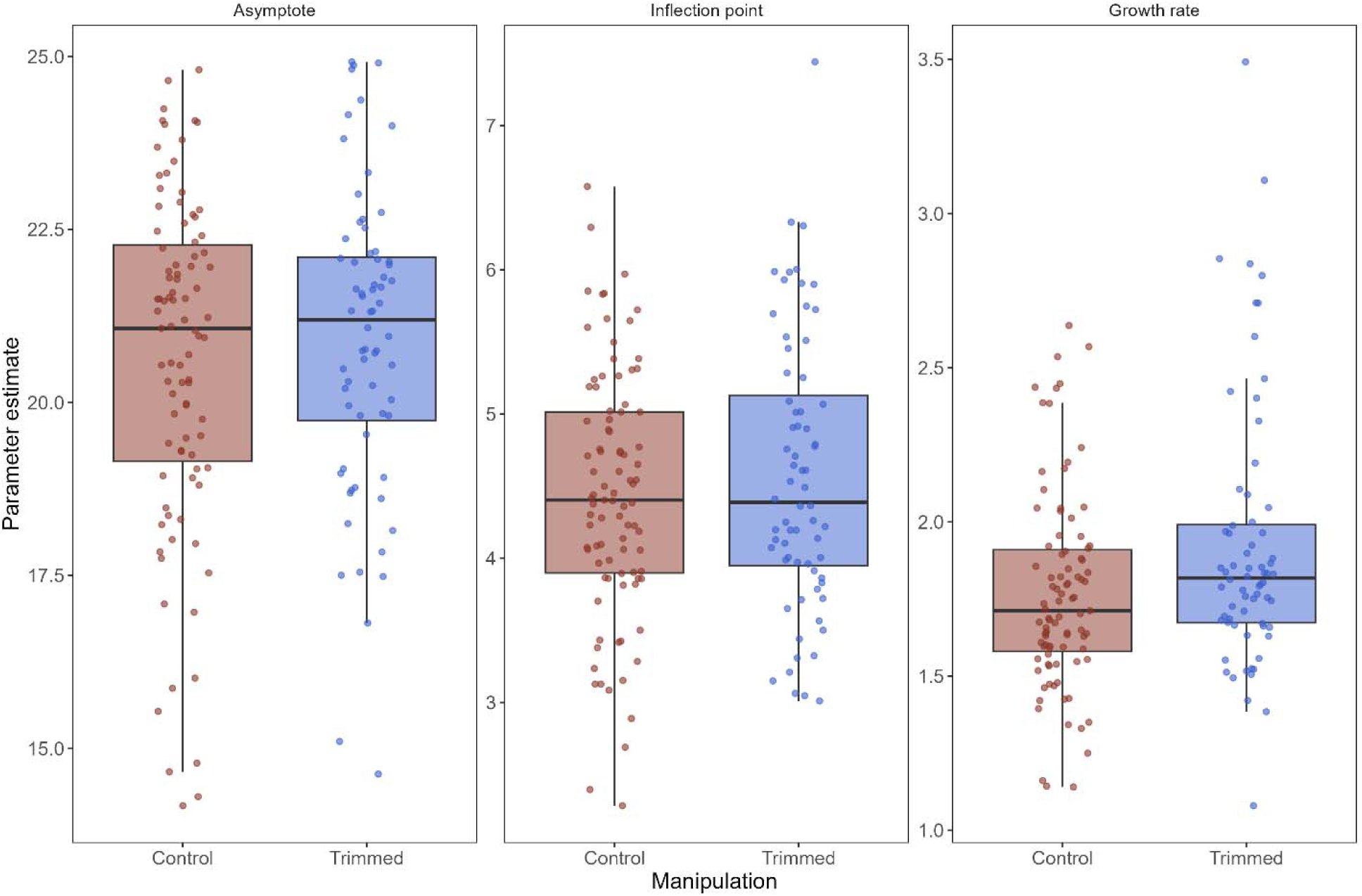
Estimates of each growth parameter (asymptote, inflection point, and growth constant) for nestlings from control and trimmed males. Each point represents the raw estimate for each nestling. Sample sizes: Control, n = 20 nests, n= 100 nestlings; trimmed, n = 15 nests, n= 86 nestlings).

**Table 7.** Parameter estimates for growth trajectories in nestling tree swallows. Estimates of fixed effects with 95% confidence intervals (CI) and *P*-values are shown. Bold indicates statistical significance. Models were run with a random effect for individual nestling identity as well as box identity (Control fathers, n = 20 nests, n= 100 nestlings; Trimmed fathers, n = 15.

| Predictor | Predictor | Estimate | 95% CI | P-value |
| --- | --- | --- | --- | --- |
| Asymptote<br>(A) | Intercept | 21.80 | 19.35, 24.26 | **<0.001** |
|  | Manipulation | 0.50 | -0.847, 1.837 | 0.471 |
|  | Brood Size | -0.30 | -0.774, 1.837 | 0.225 |
| Inflection Point<br>(I) | Intercept | 4.29 | 3.834, 4.739 | **<0.001** |
|  | Manipulation | 0.22 | 0.046, 0.397 | **0.014** |
|  | Brood Size | 0.00 | -0.088, 0.085 | 0.975 |
| Growth Rate Constant<br>(K) | Intercept | 1.88 | 1.502, 2.267 | **<0.001** |
|  | Manipulation | 0.21 | 0.063, 0.360 | **0.005** |
|  | Brood Size | -0.02 | -0.093, 0.054 | 0.599 |

nests, n= 86 nestlings).

Nestling mass was similar between small and large broods at the asymptote (P = 0.225, Table 7), inflection point (P = 0.975, Table 7), and for the growth rate (P = 0.599, Table 7).

## Discussion

We found partial support for the hypothesis that nestling provisioning rates by male tree swallows are limited by the male’s risk of overheating (i.e., the heat dissipation limit theory, Speakman & Król, 2010). Contrary to predictions, males with an enhanced capacity to dissipate heat via feather trimming did not increase their provisioning rates above the rate of untrimmed control males, nor did the two groups differ in average T_b_. Rather, male provisioning rate was largely predicted by environmental temperature and the provisioning rate of their female partners. While we anticipated that nestlings of trimmed males would have increased growth rates, our results were mixed. Taken together, our results did not provide strong support of the heat dissipation limit theory.

### Male tree swallows decreased nestling provisioning rates at high temperatures and adjusted effort to match their female partner

We anticipated that trimmed male tree swallows would maintain higher nestling provisioning rates than untrimmed control males at highest environmental temperatures, as shown previously in females (Tapper at al., 2020a). However, while we found that high environmental temperatures reduced male provisioning, trimmed males were unable (or unwilling) to increase provisioning above that of control males. A lack of response among males when provided with an increased capacity to dissipate heat suggests males may be working below their upper thermal limit. That birds may not be working maximally during parental care is an issue that has been raised previously (Williams 2018). In breeding blue tits (*Cyanistes caeruleus*), feather trimming also did not increase provisioning rates, however, trimmed males were better able to maintain body condition at the same work rate as untrimmed controls (Nord & Nilsson, 2019).

Unfortunately, we could not measure change in body condition in our study due to difficulty in recapturing male tree swallows. Although trimming did not increase male provisioning rate, the increased ability to dissipate body heat may have buffered males from the negative consequences of thermal stress. However, the effectiveness of this strategy is unclear. For example, high ambient temperatures induce oxidative stress in zebra finches (*Taeniopygia guttata)*, however, such damage was not ameliorated via trimming manipulations (Zagkle et al., 2022). Since both trimmed and untrimmed tree swallow males decreased provisioning rates at high temperatures, the physiological consequences of high environmental temperature may persist regardless of manipulation.

In general, rates of male tree swallow provisioning appear to be closely tied to those of their female partner. Previous work showed that experimental manipulation of female provisioning rate (via increased nestling begging calls) increased the provisioning rate of males, suggesting that males and females may coordinate care in a reciprocal manner (Lendvai et al., 2018). Males may also adjust investment based on the relative attractiveness of their mate, with males compensating for partners of perceived lower quality (i.e., reduced coloration; Limbourg et al., 2013) or increasing efforts when paired with high quality partners (Mahr et al., 2012). If male response is reflective of female investment or quality, then trimming manipulation may not fully account for interactions between pairs in which female quality informs male efforts (Nolazco et al., 2022).

### Male body temperature increased with provisioning rate and environmental temperature but was not affected by experimental trimming. We interpret these results with caution as our sample sizes were limited, with only four trimmed males recorded with thermal tags

Males with enhanced capacity to dissipate heat maintained a similar body temperature (T_b_) as control males, as we predicted. However, given that the experimental manipulation did not increase male work rate, our results do not support (or conflict with) the heat dissipation limit theory. However, we acknowledge that our sample sizes were limited for thermal tags (9 control and 4 trimmed males) which increases uncertainty in our effect estimates. Previous manipulations of an individual’s insulation allowed adults to maintain a lower T_b_ for the same work rate (Nord & Nilsson, 2019; Andreasson et al., 2020; Tapper et al., 2020b). Although we assume that our manipulation increased dry heat loss in trimmed male tree swallows, both trimmed and control males reduced activity at high temperatures, suggesting both groups were avoiding hyperthermia. Perhaps reduced vascularization on the male’s ventral surface (when compared with females who have a brood patch) reduced the effectiveness of our feather trimming in males and minimized the difference in heat loss between the trimmed and control groups. However, in other species males were found to have increased heat loss as a result of trimming, yet provisioning efforts were also not affected (Nord & Nilsson, 2019). An additional explanation is that male tree swallows may simply be working well below their sustained maximum (e.g., Williams, 2018) and are not at particular risk of overheating. If this were the case, then our experimental manipulation of males may be ineffective at eliciting increased parental effort.

An assumption of our test of the heat dissipation limit theory is that nest box visitation rate is an accurate metric for nestling provisioning and parental activity, however, this may not always be the case (e.g., Sarota and Williams 2019). Trimmed males may have increased investment in other activities such as self-feeding (Kacelnik, 1984; Markman, 2014) or nest defense (Swihart & Johnson, 1986; Patrick et al., 2022), which may have elevated T_b_ but would not have been detected by our PIT tag readers. Differential investment strategies between males and females may therefore complicate tests of the heat dissipation limit theory. That is, considering the different activities associated with parental investment by males and females, and the cost-benefits of each activity, heat limitations may impact aspects of male investment beyond provisioning (Harrison et al., 2009).

### Female provisioning rate was best predicted by environmental temperature and the provisioning rate of male partners

In a companion study we experimentally trimmed feathers from female tree swallows and observed an increased provisioning rate in both the trimmed females and their untrimmed male partners (Tapper et al., 2020a). Coordination between males and females may vary depending on the manipulated individual and the associated cost-benefit functions of care (Harrison et al., 2009; Bründl et al., 2025). As a result, in the current study we tested the reverse, whether females adjust their provisioning rate in response to experimental trimming of males. We found that trimming of males did not affect either male or female provisioning rate; it may be that female provisioning rate or quality drives male investment strategy more so than the reverse (Pilastro, et al., 2003; Griggio, et al., 2005; Nolazco et al., 2022). Interestingly, while male tree swallows reduced provisioning rates at the highest temperatures, females showed a slight increase. This partial compensation by females may only be possible because females can dissipate some heat via their vascularized brood patch.

### Experimental trimming of males had a modest effect on nestling growth

We found slight differences in the growth trajectories of nestlings from trimmed and untrimmed (control) males. Nestlings from both groups reached a similar asymptotic mass, however, nestlings from trimmed males grew faster, despite reaching the midpoint of their growth later than nestlings from untrimmed males. However, since males were trimmed between days 3-4 of nestling development, the slight differences in timing (for the midpoint of growth) may not reflect an actual treatment effect given the limited window of time for a true biological response. In contrast, previous work with female tree swallows found that nestlings from trimmed mothers reached a higher asymptotic mass. However, this was thought to be due to enhanced brooding capabilities of trimmed mothers (i.e., increased heat transfer to nestlings via trimming) rather than an increased provisioning rate (Tapper et al., 2020a). Since experimental trimming of males did not increase provisioning rates, a lack of difference in asymptotic mass is perhaps not surprising. Similarly, while nestlings of trimmed and untrimmed (control) fathers differed in early growth trajectories (i.e., growth constant and midpoint of growth), the magnitude of these effects was small and may not correspond to male trimming given the close overlap in timing between male trimming and the midpoint of nestling growth. Nonetheless, the differences in early growth rate suggests that male trimming, while not affecting provisioning rate, may have altered other aspect of parental care. For example, the quality of prey items (Sinkovics et al., 2021) and coordination of care between parents (Bebbington & Hatchwell, 2016) can each impact nestling growth. Overall, it remains unknown whether the small differences we detected in growth trajectories have ecological relevance.

## Conclusion

We found limited evidence that provisioning rates by male tree swallows are constrained by the male’s risk of overheating, in contrast with the heat dissipation limit theory. Although males decreased provisioning rates at high temperatures, they appeared to work routinely below a thermal limit. Importantly within a pair, males and females coordinated provisioning with environmental temperature acting as a driving factor. Our results contrast with previous experimental work on female tree swallows, in which females with an enhanced capacity to dissipate heat could maintain provisioning rates at high environmental temperatures (Tapper et al 2020a, 2020b). Overall, our study has relevance for aerial insectivores as individuals may face fitness trade-offs at high temperatures (Van de Ven et al., 2019; Cunningham et al., 2021). Different responses by males and females highlight the importance of considering sex-specific differences in thermal physiology and behavior when predicting responses to climate change.

## Acknowledgements

We thank Aleesa Schubert, Kaitlyn Baker and Sophia Luimes for assistance in the field, David Riegert for advice with data analysis, and Simon Tapper for frequent advice and discussions. Research funding was provided by a Natural Sciences and Engineering Research Council of Canada (NSERC) - Discovery Grants (awarded to GB).

## Data Accessibility

All data will be archived on the Dryad Data repository

