## Supplemental Tables 1 and 2 for "Effects of partner workload and increasing environmental temperature on nestling provisioning and body temperature in a declining aerial insectivore"

**Table S1. Model selection results for factors that explain variation in hourly provisioning rate by adult male tree swallows.** Of the 16 models run, there were three models with strong support (<2 ΔAIC_C_). Models were run with a random effect for individual bird identity (Trimmed: n=13, Control: n=18).

| **Models** | **K** | **AICc** | **ΔAICc** | **Wi** | **Acc Wi** | | **R^2^m** | **ER** |
| --- | --- | --- | --- | --- | --- | --- | --- | --- |
| **Model 1:** T_a_ + Brood Size + Provisioning Day + Relative Humidity + Wind Speed + Manipulation + Female Provisioning Rate | 8 | 8487.56 | 0.00 | 0.41 | | 0.41 | 0.358 | 1.00 |
| **Model 2:** T_a_ + Female Provisioning Rate | 5 | 8497.94 | 0.10 | 0.39 | | 0.80 | 0.362 | 1.05 |
| **Model 3**: T_a_* Manipulation + Brood Size + Provisioning Day + Female Provisioning Rate | 9 | 8489.40 | 1.84 | 0.16 | | 0.98 | 0.358 | 2.51 |
| Null Model | 3 | 9595.10 | 1107.55 | 0.00 | | 1.00 | 0.00 | >10^5^ |

Note: The null model was included for reference but did not have strong support (ΔAIC_C_ > 2). Brood Size = Brood size at hatch; Provisioning Day = Provisioning day after hatch; Relative Humidity = Average hourly relative humidity; Wind Speed = Average hourly wind speed; Female Provisioning Rate = Hourly provisioning rate of females Abbreviations are as in Table 1

**Table S2. Model selection results for factors that explain variation in hourly provisioning rate of adult female tree swallows paired with trimmed or untrimmed males.**

There were two models with strong support (<2 ΔAIC_C_). A total of 16 candidate models were run with a random effect for individual bird identity (females partnered with Trimmed males, n= 13; females partnered with Control males, n= 17).

| **Models** | **K** | **AICc** | **ΔAICc** | **Wi** | **Acc Wi** | | **R^2^m** | **ER** |
| --- | --- | --- | --- | --- | --- | --- | --- | --- |
| **Model 1:** T_a_ + Manipulation + Male Provisioning Rate | 6 | 1630.70 | 0.00 | 0.67 | | 0.67 | 0.336 | 1.00 |
| **Model 2:** T_a_ + Brood Size + Provisioning Day + Relative Humidity + Wind Speed + Manipulation + Male Provisioning Rate | 10 | 1632.10 | 1.39 | 0.33 | | 0.99 | 0.363 | 2.01 |
| Null Model | 3 | 2336.25 | 705.55 | 0.00 | | 1.00 | 0.00 | >10^5^ |

Note: The null model was included for reference but did not have strong support (ΔAIC_C_ > 2). Abbreviations are as in Table S1
